# Parkinson’s Disease Pathology is Directly Correlated to SIRT3 in Human Subjects and Animal Models: Implications for AAV.SIRT3-myc as a Disease-Modifying Therapy

**DOI:** 10.1101/2023.06.23.546104

**Authors:** Dennison Trinh, Ahmad R. Israwi, Harsimar Brar, Jose E.A. Villafuerte, Ruella Laylo, Humaiyra Patel, Shaumia Sinnathurai, Kiran Reehal, Alyssa Shi, Vayisnavei Gnanamanogaran, Natalie Garabedian, Drake Thrasher, Philippe P. Monnier, Laura A. Volpicelli-Daley, Joanne E. Nash

## Abstract

Degeneration of the dopaminergic nigro-striatal pathway and presence of Lewy bodies are pathological hallmarks of Parkinson’s disease (PD). Postmortem studies in human tissue have also demonstrated that a decline in mitochondrial number and function is also central to PD pathology. Sirtuin 3 (SIRT3) is a mitochondrial protein deacetylase which has been linked with longevity and cytoprotective effects. SIRT3 serves as a metabolic sensor and regulates mitochondrial homeostasis and oxidative stress, which likely stabilises telomere integrity, delaying senescence. Previously, we have shown that over-expression of SIRT3 rescues motor function and prevents degeneration of dopaminergic neurons in a virally over-expressing mutant (A53T)-α-synuclein model of PD. In the present study, we show that in the substantia nigra *pars compacta* (SNc) of human subjects, SIRT3 levels are negatively correlated with age (p<0.05, R=0.6539). In the hippocampus, there was no correlation between SIRT3 levels and age. In human subjects with PD, SIRT3 was reduced by 56.8±15.5% and 34.0±5.6% in the SNc and hippocampus respectively regardless of age. Given that age is the primary risk factor for PD, this finding suggests that reduced SIRT3 may be a causative factor contributing to PD pathology. Next in human subjects with PD, we measured whether there was a correlation between the amount of aggregated α-synuclein and SIRT3 levels by measuring immunofluorescence of phosphorylated α-synuclein (p-syn), which is a marker for Lewy bodies. Interestingly, in the hippocampus, but not SNc, there was a positive correlation between SIRT3 and p-syn levels, despite p-syn being reduced compared to control. Next using an α-synuclein seeding rat model of PD, we assessed the disease-modifying effects of viral-mediated SIRT3 infusion. Six months following infusion of α-synuclein pre-formed fibrils (PFF) into the SNc, there was 38.8±4.5% loss of TH-positive neurons, impaired striatal dopamine metabolism and pathological α-synuclein throughout the brain. Phosphorylated-α-synuclein immunoreactivity was present in the SNc, olfactory tubercle, striatum, amygdala, hippocampus and motor cortex. In PD subjects, synuclein positive aggregates have also been reported in these brain regions. In PFF rats, infusion of rAAV1.SIRT3-myc in the SNc reduced abundance of α-synuclein inclusions in the SNc by 30.1±18.5% which was not seen when deacetylation deficient SIRT3^H248Y^ was transduced. This demonstrates the importance of SIRT3deacetylation in reducing α-synuclein aggregation. However, while SIRT3 transduction reduced aggregation in the SNc, it had no significant effect on phosphorylated-α-synuclein levels in other brain regions. These studies confirm that SIRT3 is directly correlated with senescence and aging in humans. We also provide evidence that reduced SIRT3 contributes to the pathology of clinical PD. Finally, by showing that over-expression of SIRT3 prevents α-synuclein aggregation through de-acetylation-dependent mechanisms, we further validate AAV1.SIRT3-myc as a potential disease-modifying therapy for PD.

## Introduction

Parkinson’s disease (PD) is the most prevalent neurodegenerative movement disorder worldwide, affecting approximately ten million individuals globally. The cardinal symptoms of PD are bradykinesia, resting tremor and muscle rigidity, as well as non-motor symptoms, such as autonomic dysfunction and cognitive decline. Presently only symptomatic treatments are available with no impact on disease progression. The prevalence of PD is predicted to double in the next ten years (Ou et al., 2021), therefore, the need for disease-modifying agents is imperative.

PD is confirmed post-mortem by degeneration of dopaminergic neurons in the substantia nigra pars compacta (SNc) and the presence of Lewy bodies, which are spherical, intracellular inclusions comprised of mis-folded ubiquitinated and phosphorylated α-synuclein, p62, organelles and lipids (Duffy and Tennyson, 1965; Mahul-Mellier et al., 2020; Shahmoradian et al., 2019). Mitochondria regulate critical processes such as ATP production (Bertram et al., 2006), calcium and cellular homeostasis (Romero-Garcia and Prado-Garcia, 2019), oxidative stress (Zhao et al., 2019) and proteostasis (Moehle et al., 2019), all of which have been linked with neurodegeneration in cell and animal models of PD (Ferreon and Deniz, 2007). Mitochondria are especially important in dopaminergic neurons, as these neurons have higher energetic demands and are more sensitive to stress compared to other neuronal cell types (Pacelli et al., 2015; Surmeier and Bevan, 2003). Post mortem studies in human tissue show that aberrant mitochondria function is central to PD pathology, with increased mitochondrial DNA mutations and oxidative stress, as well as reduced ATP and electron transport function being present in both idiopathic and familial cases (Gu et al., 1998; Trinh et al., 2020). Importantly, inhibition of mitochondria complex I of the electron transport chain in humans and animals, e.g. using mitochondrial toxins such as MPTP (Bezard et al., 1997) and rotenone (Cannon et al., 2009) or through genetic manipulation (Gonzalez-Rodriguez et al., 2021) is sufficient to induce cardinal symptoms of Parkinson’s disease, as well as degeneration of the substantia nigra pars compacta, regardless of the presence or absence of α-synuclein aggregates.

Pre-clinical studies have shown that enhancement of mitochondrial health has disease-modifying effects (Bido et al., 2017; Park et al., 2020; Shi et al., 2017; Zhou et al., 2019). Early clinical trials involving mitochondrial enhancers, NAD+ (Perez et al., 2021), creatine (Mo et al., 2017) and co-enzyme Q10 (Zhu et al., 2017) failed, likely because of small cohort numbers and low bioavailability (Chen-Plotkin et al., 2018; Dawson and Dawson, 2019; Paolini Paoletti et al., 2020). However, more recently, enhancement of mitochondrial health has also shown promise in Phase I clinical trials. In 2022, nicotinamide riboside (NR) was shown to have mild symptomatic benefits in PD subjects (Brakedal et al., 2022). NR is the precursor to NAD+ which regulates stress responses through the regulation of mitochondrial homeostasis and genomic stability. Importantly, NAD+ decreases with age, and is decreased further in PD subjects (Massudi et al., 2012; Mischley et al., 2023). It is currently unknown whether low NAD+ levels are a rate limiting factor in the disease-modifying ability of NR.

Sirtuin 3 (SIRT3) is a NAD+–dependent protein deacetylase localised to the mitochondria. SIRT3 is associated with cytoprotection and longevity in organisms ranging from yeast and c-elegans to humans. SIRT3 serves as a metabolic sensor, activated by metabolic and cellular allostasis and increased availability of NAD+. Many of the protective effects of NAD+ are mediated through the de-acetylase effects SIRT3 (van de Ven et al., 2017). Indeed deacetylation-deficient mutations in SIRT3, such as the H248Y result in a loss of the cytoprotective properties of SIRT3 (Gleave et al., 2017)Importantly, as well as requiring NAD+ for activation, SIRT3 activation also restores NAD+ levels, thus SIRT3 may be more attractive as a mitochondrial enhancing target (Fulco et al., 2008; Xie et al., 2017). Therefore, elevation of SIRT3 may represent a mechanism for enhancing mitochondria function, which also has the ability to maintain NAD+ levels. In neuronal stem cells, SIRT3 function declines with age, and age is the most predominant risk factor for PD. This indirectly suggests that declining SIRT3 levels may be causative in the pathology of PD (Joseph et al., 2012; Ou et al., 2021). At the molecular level, SIRT3 enhances mitochondrial health by stabilising mitochondrial membrane processes including the electron transport chain (Gleave et al., 2017) to reduce metabolic energy demands (Fu et al., 2022; Hirschey et al., 2010; Zhang et al., 2016) and oxidative stress (Chen et al., 2011; Iwahara et al., 2012; Li et al., 2015; Tao et al., 2010; Tseng et al., 2013; Zhang et al., 2016), making it an excellent candidate as a disease-modifying therapy for PD. Our previous studies showed that in a rat model of PD, over-expression of SIRT3 using AAV1.SIRT3-myc, is neurorestorative, causing motor function recovery, preventing neurodegeneration, and reversing abnormalities in striatal dopamine metabolism (Gleave et al., 2017). However, no single animal model emulates the pathology of clinical PD. This is partly because of the inherent problems with animal and cell models, also because PD is a heterogenous syndrome caused by a combination of genetic and environmental risk factors. Thus, pre-clinically, potential disease-modifying agents should be validated using multiple models.

In this study, we firstly link reduced SIRT3 to the pathology of PD in human subjects. Next, we evaluated the efficacy of AAV1.SIRT3-myc as a disease-modifying agent in an animal model that recapitulates the spread of α-synuclein fibrils through the brain, the α-synuclein pre-formed fibril (PFF) rat. We demonstrate that in the PFF rat, SIRT3 over-expression reduces α-synuclein aggregation *in vivo*.

## Results

### SIRT3 is reduced in human subjects with Parkinson’s disease

Human subjects with PD were assessed post-mortem for SIRT3, acetyl-lysine, and phosphorylated Ser129 α-synuclein (p-α-syn) protein levels using SDS-PAGE followed by Western blot and compared to age-matched controls (Fig. 1A-C). As has been shown previously, p-α-syn which is an indicator of aggregated α-synuclein Lewy pathology was increased relative to control subjects by 96.5±36.4% and 85.9±23.3% in the SNc and hippocampus of PD subjects respectively. Conversely, SIRT3 was reduced in PD subjects relative to age-matched controls by 56.8±15.5% and 34.0±5.6% in the SNc and hippocampus respectively. Since SIRT3 is a mitochondrial protein deacetylase, cellular acetyl-lysine levels were measured as the total amount of acetylated proteins across all molecular weights, however, there was no significant difference in acetyl-lysine levels between PD and control subjects (Fig. 1B, 1C). In the SNc of control patients there was a strong inverse correlation between age and SIRT3 levels (R=0.6539), whereby SIRT3 levels declined with increasing age (Fig. 2Ai). No correlation was observed between acetyl-lysine and age, p-α-syn and duration, or SIRT3 and p-α-syn in the SNc (Fig. 2Aii-iv). Furthermore, no correlation was observed between SIRT3 and age, acetyl-lysine and age, or p-α-syn and duration in the hippocampus (Fig. 2Bi-iii). Lastly, in the hippocampus of PD subjects there was a positive correlation between p-α-syn and SIRT3 levels which was not observed in control subjects (R=0.6344) (Fig. 2Biv).

**Figure 1.**
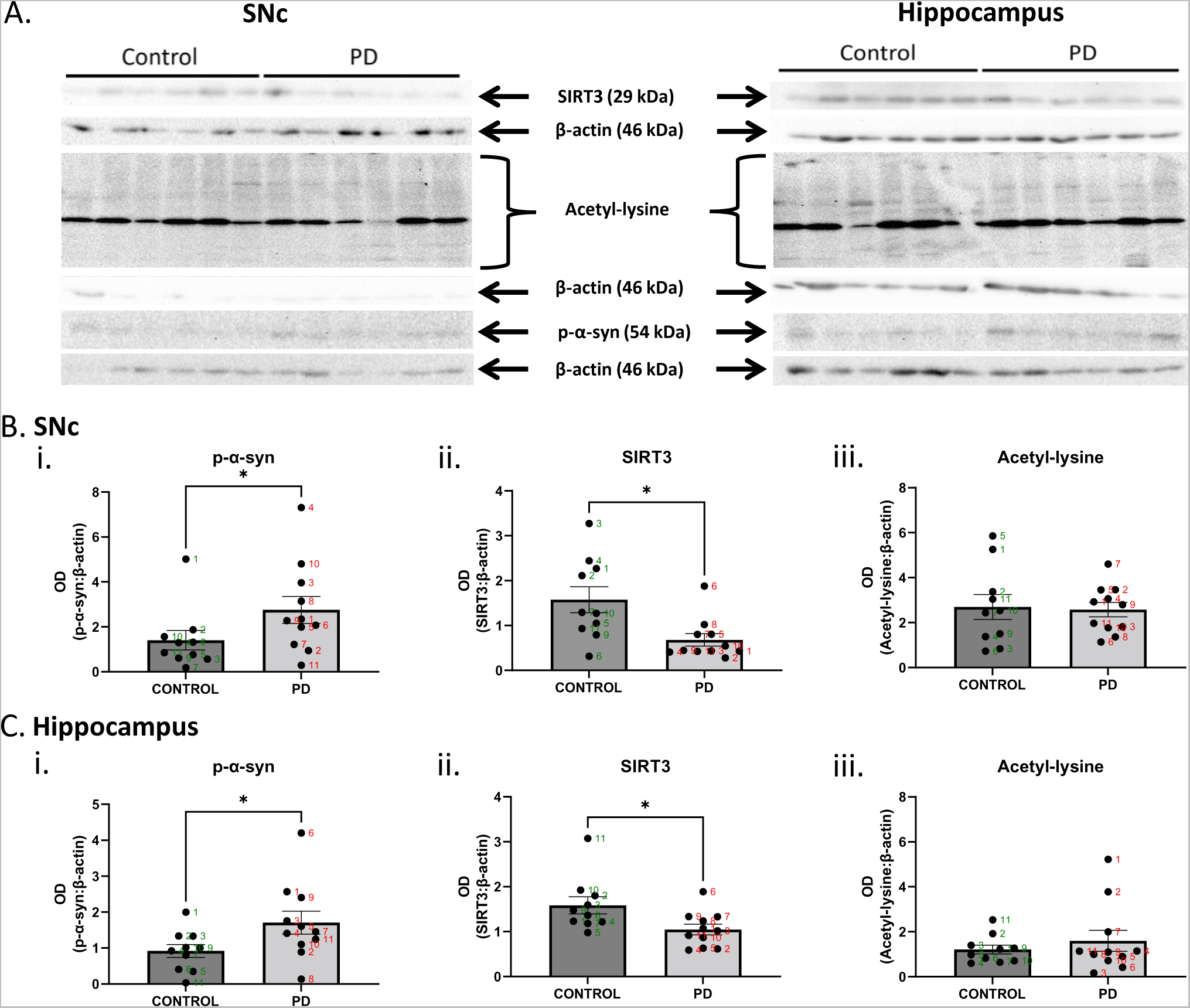
In human post-mortem brain tissue from PD subjects, SIRT3 is decreased and p-*α*-syn is increased in the SNc and hippocampus. **A.** SDS-PAGE followed by Western Blot was performed on homogenized tissue from the SNc and hippocampus using antibodies against p-α-syn, SIRT3, acetyl-lysine, and β-actin (loading control). **Bi**. p-α-synuclein, **Bii.** SIRT3, **Biii.** Acetyl-l-lysine levels in the SNc quantified using optical density (mean ± SEM) as a ratio of β-actin. **Ci**. p-α-synuclein, **Cii.** SIRT3, **Ciii.** Acetyl-lysine levels in the hippocampus quantified using optical density (mean ± SEM) as a ratio of β-actin. **D.** Mean protein levels ± SEM in the SNc for **Di.** SIRT3 and **Dii.** Acetyl-lysine relative to age (years). Numbers in B–D correspond to individual subjects from each group, with subject 9 and 10 being different for the SNc and hippocampus tissue for the control group. n = 8-11 per group. Mann-Whitney U-Test. * p<0.05

**Figure 2.**
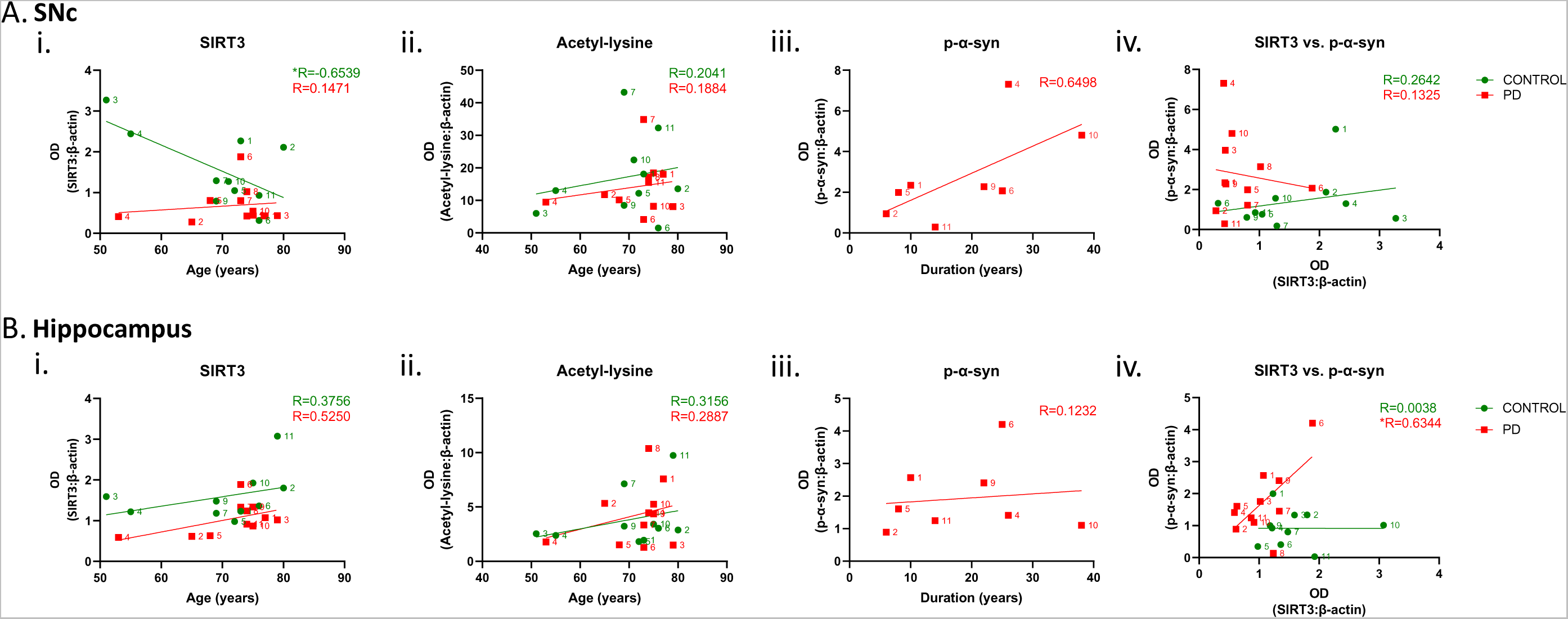
In human post-mortem brain tissue from control subjects, SIRT3 declines with age in the SNc while in PD subjects, SIRT3 increases with p-α-syn in the hippocampus. **Ai.** SIRT3 and **Aii.** Acetyl-lysine relative to age (years), **Aiii.** p-α-syn compared to duration of PD, and **Aiv.** p-α-syn compared to SIRT3 in the SNc with linear regression analysis and R values shown. **Bi.** SIRT3 and **Bii.** Acetyl-lysine relative to age (years), **Biii.** p-α-syn compared to duration of PD, and **Biv.** p-α-syn compared to SIRT3 in the hippocampus with linear regression analysis and R values shown. Numbers in B–D correspond to individual subjects from each group, with subject 9 and 10 being different for the SNc and hippocampus tissue for the control group. n = 8-11 per group. Pearson Correlation Test. * p<0.05

### Intra-nigral-infusion of PFFs causes degeneration of dopaminergic neurons and alters striatal dopamine metabolism but has no effect on forelimb use

Following nigral infusion of PFFs and AAV1.SIRT3-myc, forelimb asymmetry was assessed at 1, 3 and 6month time points and compared to baseline. Infusion of PFFs alone caused no significant change in forelimb use compared to control (monomeric α-synuclein (mono+EV)) at any of the time points assessed (Fig. 3). Nigral infusion of both AAV1.SIRT3-myc (PFF+SIRT3) and the deacetylation-deficient SIRT3 mutant (PFF+SIRT3^H248Y^) had no significant effect on forelimb use compared to either the mono+EV or PFF+EV group.

**Figure 3.**
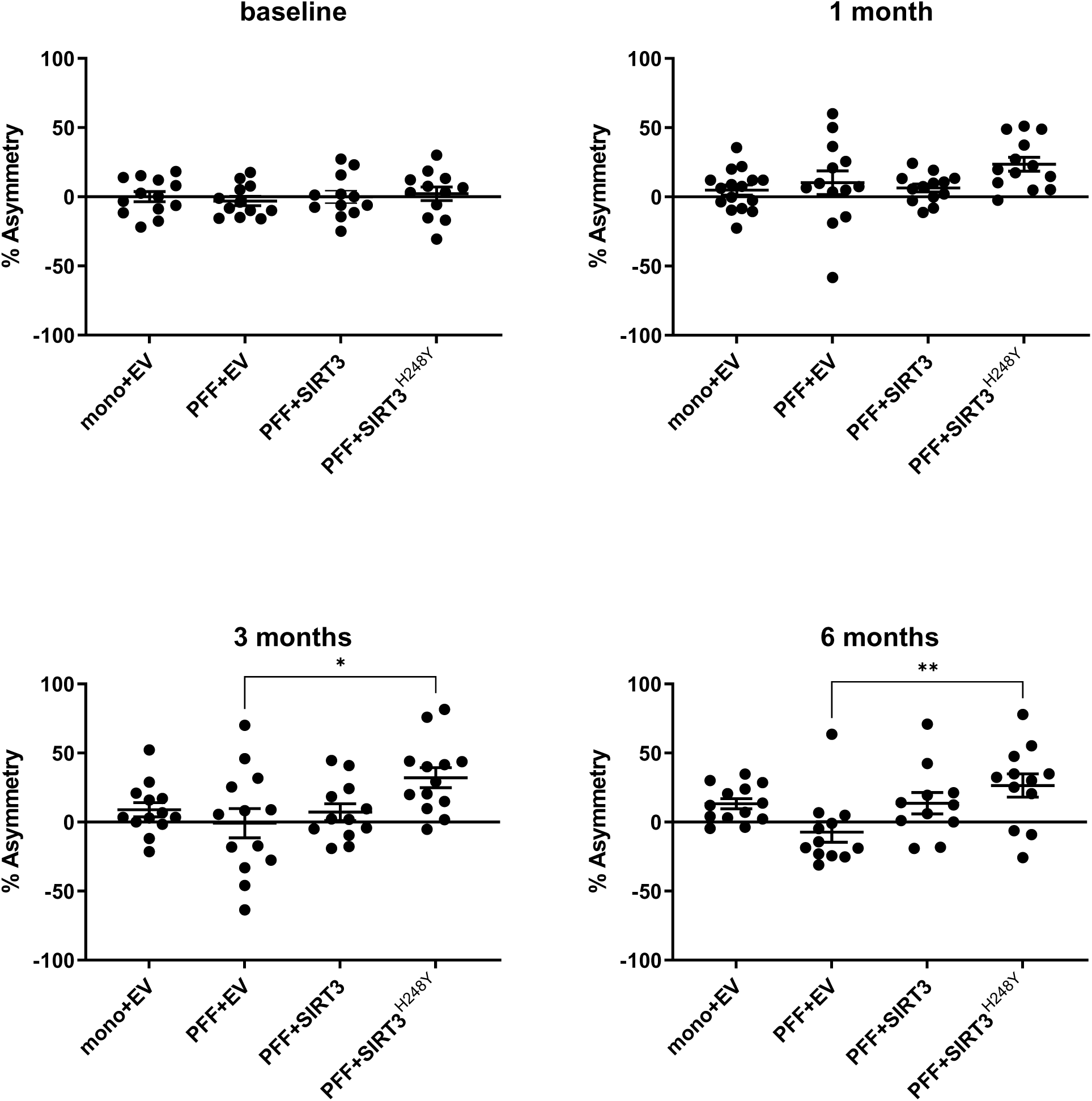
There is no effect of PFFs and AAV1.SIRT3-myc on motor coordination in rats. Infusion of PFFs into the SNc had no significant effect on forelimb asymmetry at 1, 3, or 6 months (n = 11-16 per group). One-way ANOVA test with Tukey’s multiple comparisons *post-hoc*. * p<0.05, ** p<0.01.

Next, we quantified the number of dopaminergic neurons in the SNc. Six months post PFF / AAV.SIRT3 delivery, animals were euthanised, brains removed, post-fixed in 4% paraformaldehyde, then processed for stereological quantitation of dopaminergic neurons in the SNc. Following PFF infusion (PFF+EV) there was significant loss of TH-positive (dopaminergic) neurons in the ipsilateral SNc (38.8±4.52%) compared to control (mono+ EV) (Fig. 4). In the SNc of the PFF+EV group on the side contralateral to PFF infusion, there was no significant decrease in dopamine cell number compared to control (mono+EV) (Fig. 4) which was expected since a unilateral infusion was utilised for this study. AAV.SIRT3-myc had no effect on the number of TH-positive cells. There was no significant difference between the PFF+SIRT3 injected and PFF+EV groups (1512±398.4 and 2159±196.6 TH-positive cells respectively). In the (PFF+SIRT3 group, the decrease in TH-positive neurons was comparable to that observed in the PFF+EV group (57.12±15.61%) compared to control (mono+EV) (Fig 4). As was the case with wild-type SIRT3, the enzymatically inactive (deacetylation-deficient SIRT3 mutant SIRT3^H248Y^) did not rescue neuronal loss caused by PFF in the ipsilateral side, but unexpectedly we observed an increase in dopaminergic neurons on the contralateral side of rats exposed to PFF and the mutated protein (Fig 4). This may be due to an unknown compensatory mechanism in the contralateral hemisphere in response to the elevated inactive SIRT3^H248Y^ expression in the ipsilateral hesmiphere. Irrespective of this, the principal conclusion from these data is that PFF depletes dopaminergic cells in the SNc surrounding the site of injection, and SIRT3 does not prevent that loss (Fig 4).

**Figure 4.**
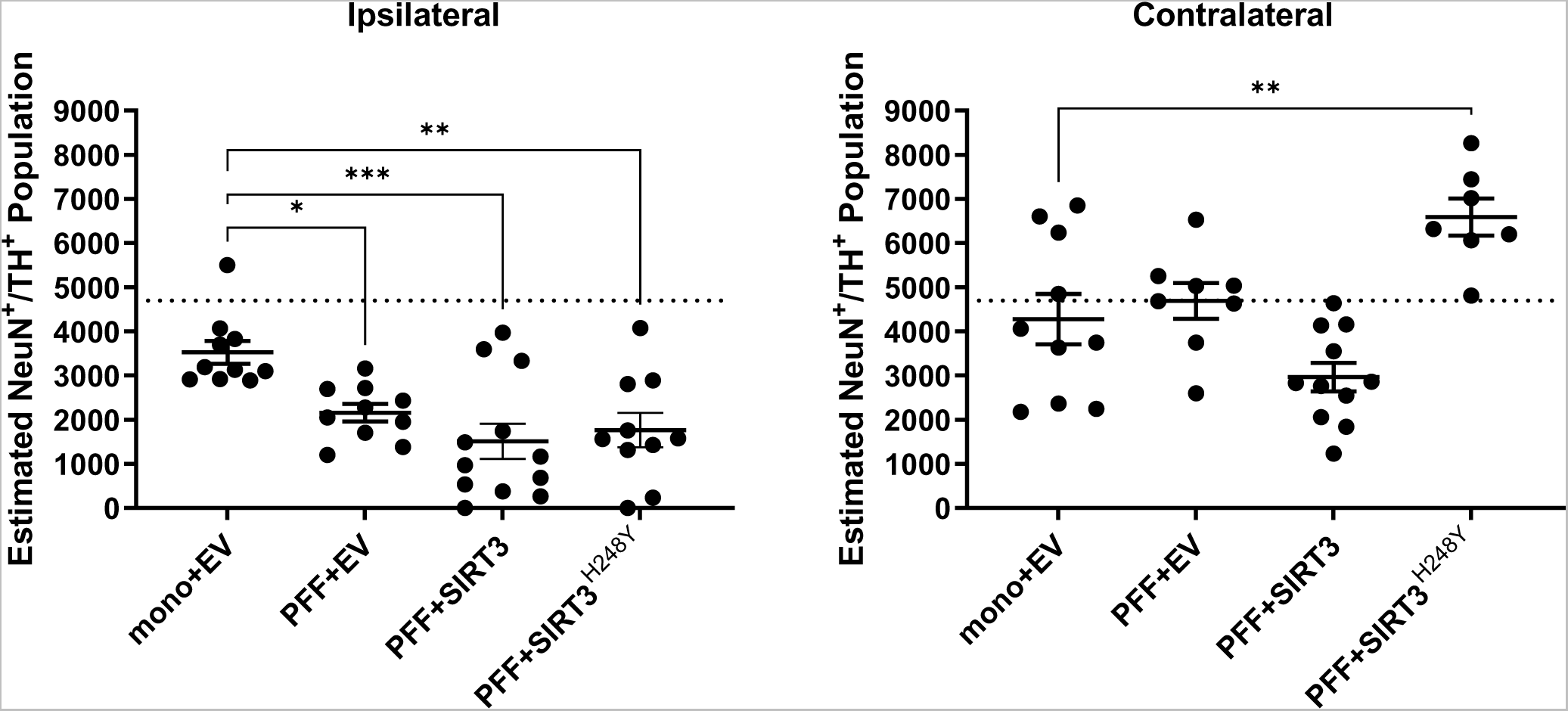
Effect of PFFs and AAV1.SIRT3-myc on dopaminergic cell number. Dopaminergic cell number was quantified in the SNc using quantitative stereology with NeuN (neuronal marker) and TH (dopamine marker). Decreased dopaminergic cell number was observed in the ipsilateral hemisphere in the PFF rat model, with no protective effects following transduction of SIRT3-myc (n = 8-12 per group). The dotted line represents the number of dopaminergic neurons in the SNc of naive animals. One-way ANOVA test with Bonferroni’s multiple comparisons *post-hoc*. * p<0.05, ** p<0.01, *** p<0.001.

Six months following nigral infusion of PFFs, dopamine and dopamine metabolites, 3,4-dihydroxyphenylacetic acid (DOPAC), 3-methoxytyramine (3-MT), and homovanillic acid (HVA) were analysed in the striatum using high performance liquid chromatography (HPLC). Striatal dopamine or dopamine metabolites were quantified relative to dopamine as a percentage of levels found in the contralateral uninjected hemisphere. In the PFF+EV group, there was a 69.0±10.3% decrease in striatal dopamine compared to control (mono+EV) (Fig. 5). In contrast, DOPAC, 3-MT and HVA levels were elevated by 40.2±3.5%, 59.4±1.4% and 42.9±2.% respectively (Fig. 5). This pattern of reduced striatal dopamine and increased striatal dopamine metabolites is indicative of dysfunctional striatal dopamine metabolism. This was expected in the PFF model given the loss of dopaminergic neurons observed in the ipsilateral SNc (Fig.4). Following AAV.SIRT3 and SIRT3^H248Y^ infusion there was no effect on dopamine cell number or dopamine metabolism (Fig 5). Thus, while PFF does not affect behaviour, it depletes ipsilateral substantia nigra dopaminergic neurons, resulting in the expected reduction or increase in striatal dopamine or dopamine metabolites, respectively, and SIRT3 does not alter any of these parameters.

**Figure 5.**
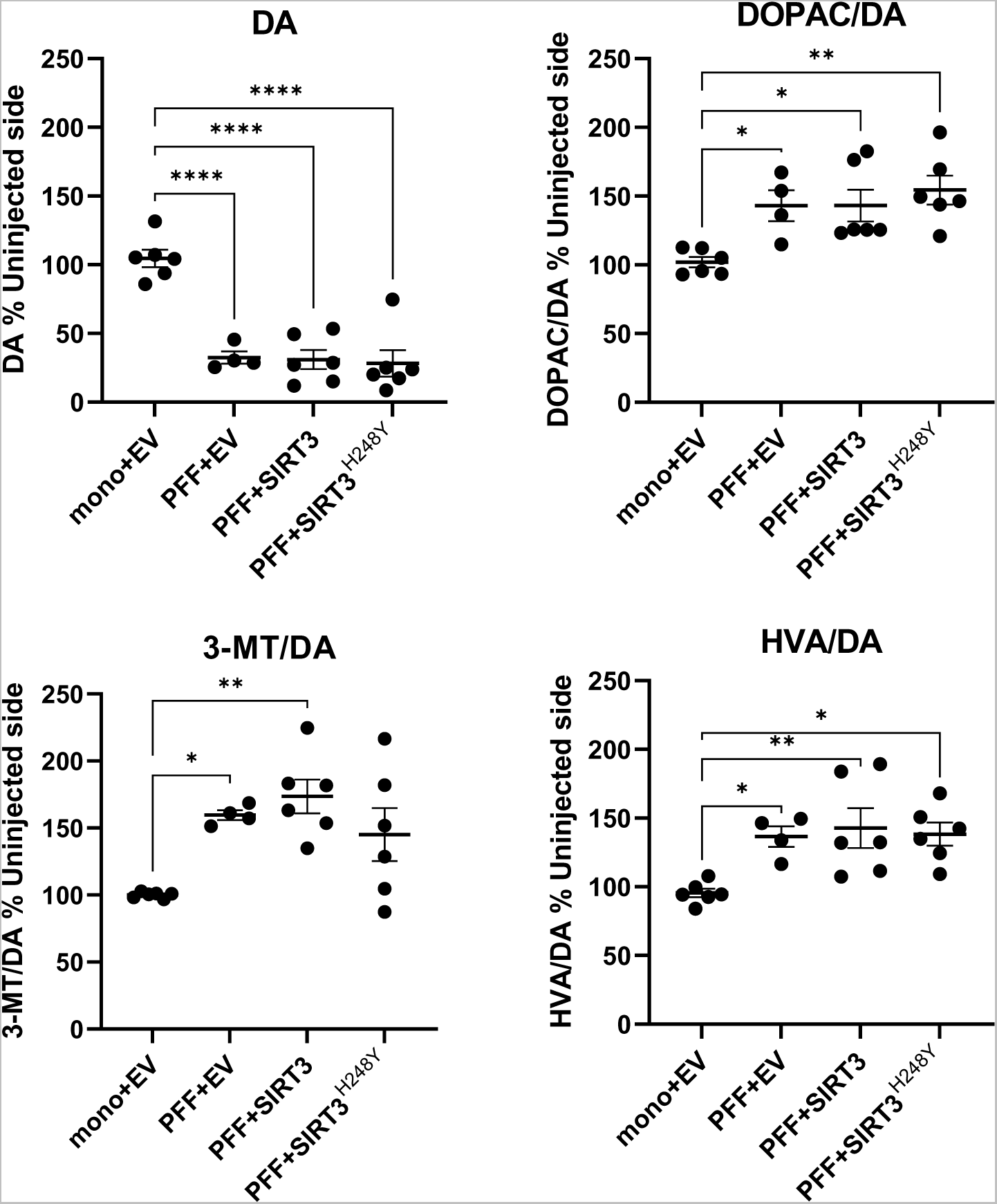
Effect of PFFs and AAV1.SIRT3-myc on striatal dopamine metabolism. Analysis of striatal dopamine (DA) and metabolites, 3,4-dihydroxyphenoylacetic acid (DOPAC), 3-methoxytyramine (3-MT), and homovanillic acid (HVA) using HPLC. Following nigral infusion of PFFs, there was a significant decrease in striatal levels of dopamine and dopamine metabolites, which was not altered following SIRT3-myc transduction (n = 4-6 per group). One-way ANOVA test with Bonferroni’s multiple comparisons *post-hoc*. * p<0.05, ** p<0.01, *** p<0.001, **** p<0.0001.

### SIRT3 decreases the spread of phosphorylated-α-synuclein in the brain

Phosphorylated-α-synuclein (p-α-syn) is a marker for pathological aggregated α-synuclein, a component of Lewy bodies that are a hallmark of PD. Thus, immunofluorescence labeling was performed against phosphorylated Ser129 α-synuclein (p-α-syn). Six months following nigral infusion of PFFs, p-α-syn positive aggregates were apparent in the SNc, striatum, hippocampus, motor cortex, olfactory tubercule and amygdala (Fig. 6–8). In human subjects with PD all of these brain nuclei show Lewy body pathology (Braak et al., 2004; Braak et al., 2003).

**Figure 6.**
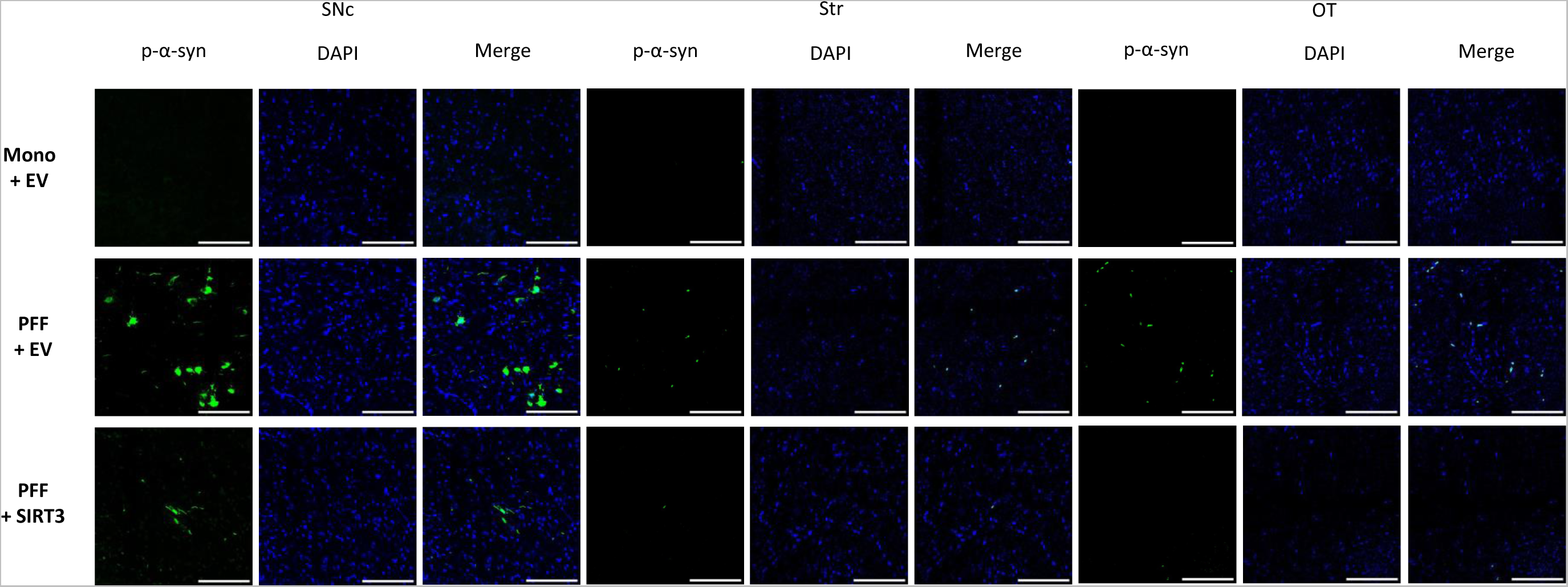
Pathological deposits of α-synuclein can be visualized using antibodies against phosphorylated Ser129 α-synuclein (p-α-syn). Representative images of p-α-synuclein immunofluorescence labelling from substantia nigra pars compacta, striatum, and olfactory tubercule brain regions affected in PD according to those proposed in Braak staging taken from the ipsilateral hemisphere following infusion of PFFs into the right SNc. Scale bar = 100 μm

### Transduction of SIRT3-myc decreased the spread of phosphorylated-α-synuclein in the brain

In the PFF rats, immunostaining and heatmap representation suggested that SIRT3 reduced p-α-syn abundance in the SNc, but not other brain regions (Fig 6, 7). Quantification confirmed that there was a significant effect of AAV1.SIRT3-myc in the PFF+SIRT3 group, where there was a significant (30.1±18.5%) reduction in p-α-syn abundance in the SNc compared to the PFF+EV group (P<0.05, Fig 6–8). In contrast, there was no significant reduction in p-α-syn abundance in the SNc with over-expression of the deacetylation-deficient SIRT3 mutant (PFF+SIRT3^H248Y^) compared to PFF+EV (P>0.05, Fig 6 and 8). In all other brain regions analysed, SIRT3 over-expression, whether SIRT3-myc or SIRT3^H248Y^, did not reach significance in altering p-α-syn levels, compared to PFF-EV (all P>0.05, Fig 6 and 8). While there was no significant effect of SIRT3 on p-α-syn levels in the striatum, OT, amygdala, motor cortex or hippocampus, this may have been due to the low number of replicates in these brain regions.

**Figure 7.**
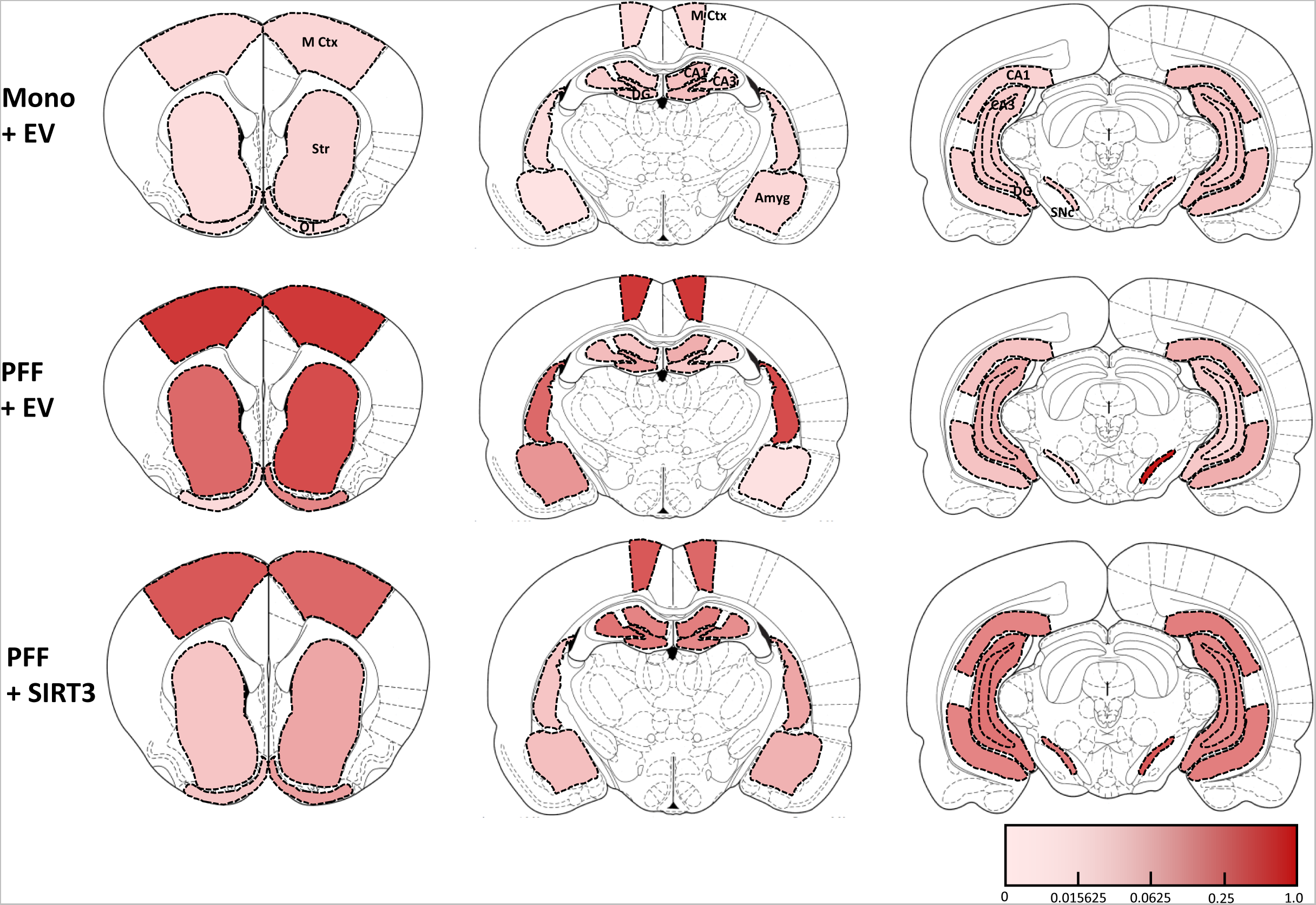
Infusion of PFFs into the SNc induces a model of PD that follows a Braak staging pattern. Heat map showing p-α-syn abundance in select regions throughout the brain. Brain sections are 1.70, −2.80, and −6.04 mm relative to Bregma respectively. **Abbreviations:** SNc – substantia nigra pars compacta, Str – striatum, M Ctx – motor cortex, Amyg – amygdala, CA1 – hippocampal CA1, CA3 – hippocampal CA3, DG – dentate gyrus.

**Figure 8.**
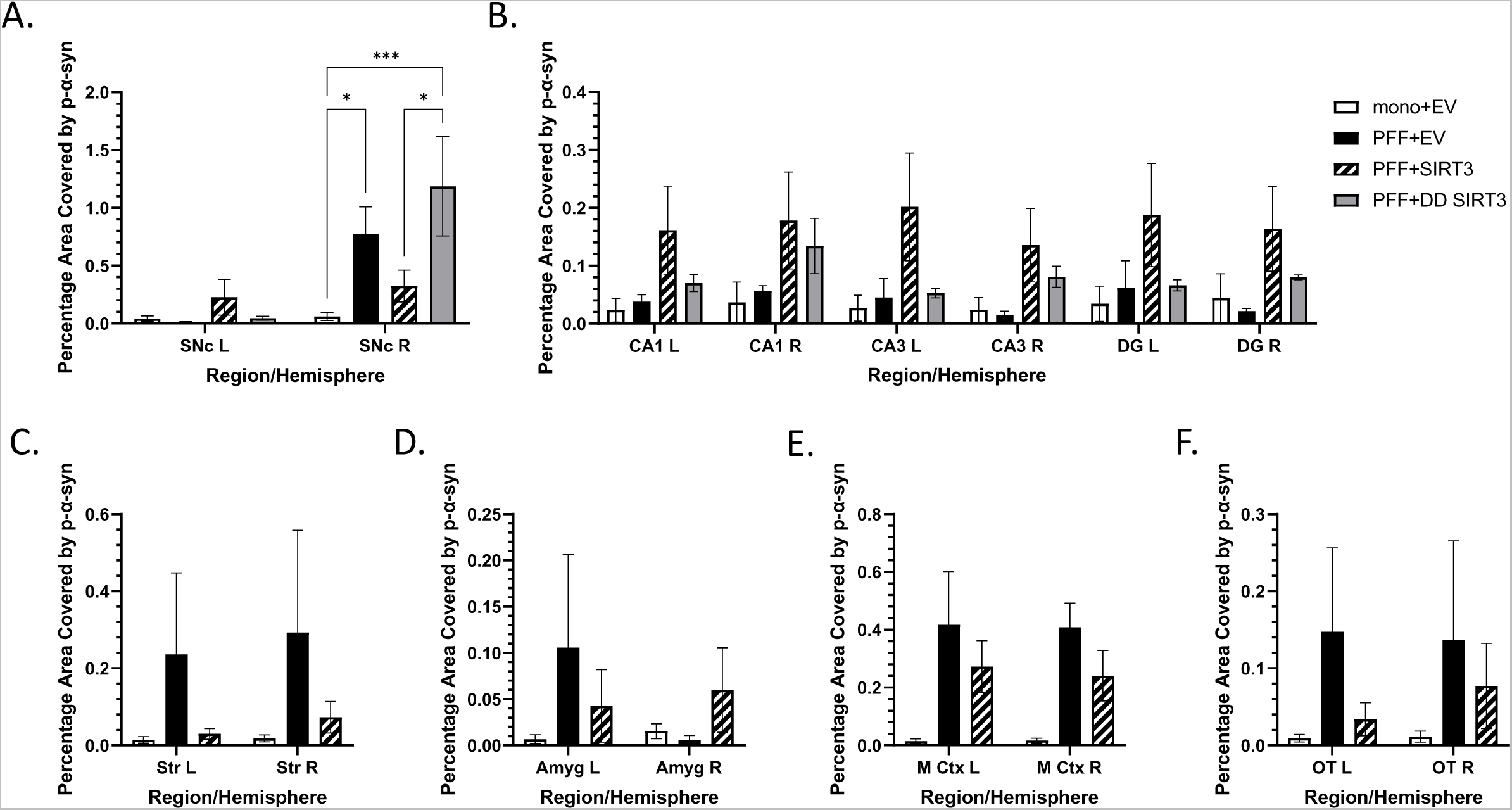
In the nigral PFF rat model of PD, SIRT3 overexpression reduces pathological *α*-synuclein aggregation in the SNc. Quantitative analysis of phosphorylated Ser129 α-synuclein (p-α-syn) abundance in the rat brain **A.** substantia nigra, **B.** hippocampus, **C.** striatum, **D.** amygdala, **E.** motor cortex, and **F.** olfactory tubercule, regions previously shown to be affected in PD. Significantly higher levels of p-α-syn fibrils and Lewy bodies were observed in the PFF+DD SIRT3 group in the SNc compared to the PFF+SIRT3 group, suggesting that SIRT3-myc over-expression reduces the spread of p-α-syn following PFF infusion in the SNc. n = 2-4 per group. **Abbreviations:** SNc – substantia nigra pars compacta, Str – striatum, M Ctx – motor cortex, Amyg – amygdala, CA1 – hippocampal CA1, CA3 – hippocampal CA3, DG – dentate gyrus. Two-way ANOVA test with Bonferroni’s multiple comparisons *post-hoc*. * p<0.05, ** p<0.01, *** p<0.001

## Discussion

The cytoprotective and longevity enhancing abilities of SIRT3 have been documented in many organisms ranging from yeast to humans (Bellizzi et al., 2005; Kong et al., 2010; Onyango et al., 2002; Shi et al., 2005). The beneficial effects of SIRT3 are mediated through its multi-faceted role in pathways critical to the regulation of ATP production (Bertram et al., 2006), calcium and cellular homeostasis (Romero-Garcia and Prado-Garcia, 2019), oxidative stress (Zhao et al., 2019) and proteostasis (Moehle et al., 2019). We have previously demonstrated that intra-nigral infusion of AAV1.SIRT3-myc has both neuroprotective and neurorestorative effects in rodent and cell models of PD (Gleave et al., 2017). In the previous study, we showed that in the virally transduced A53T mutant α-synuclein rat model of PD, AAV1.SIRT3-myc transduction provides functional recovery of motor abnormalities and reverses cellular stress to prevent neurodegeneration (Gleave et al., 2017). We have also shown that these disease-modifying effects are mediated through SIRT3 targeting the mitochondria to reduce acetylated protein levels, which stabilises oxidative respiration, reducing the metabolic demand on dopaminergic cells, and thus making them less susceptible to cell stress (Gleave et al., 2017). To further validate SIRT3 as a disease-modifying target, we assessed whether SIRT3 is linked to the pathogenesis of clinical PD, and also whether over-expression of SIRT3 is protective in a model of PD which shows spread of aggregated α-synuclein, the PFF rat model.

### Reduced SIRT3 may contribute to the pathology of clinical Parkinson’s disease

Previous studies in animal and cell models of PD have indicated that reduced SIRT3 may contribute to PD pathology. In SIRT3 KO mice, MPTP-induced loss of dopaminergic neurons is enhanced compared to wild-type mice, suggesting that endogenous SIRT3 prevents degeneration of dopaminergic neurons. Furthermore, in heterologous cells and *in vivo*, following viral-mediated transduction of mutant α-synuclein, α-synuclein aggregates invade mitochondria to decrease SIRT3 (Park et al., 2020). Finally, in DJ-1 KO mice, there is an age-related loss of SIRT3 function, which is linked with increased oxidative stress and subsequent dopaminergic cell death in the SNc (Shi et al., 2017). In this study, to determine whether SIRT3 levels are correlated with clinical PD, we measured endogenous levels of SIRT3 in the SNc and hippocampus of human subjects with PD and compared them to age-match controls. Regression analysis showed that in control subjects, there was an inverse relationship between SIRT3 and age, whereby SIRT3 declined with increasing age. Conversely in clinical PD, SIRT3 was reduced in all subjects. Since age is the primary risk factor for PD, and SIRT3 both declines with age, and is uniformly low in PD, this suggests that lack of SIRT3 may contribute directly to PD pathology. Interestingly, in the hippocampus of clinical PD, as p-α-syn levels increased, SIRT3 levels also increased though not to the levels observed in the control subjects. This may be linked to increased neurogenesis in the hippocampus, which overall may elevate SIRT3 levels in the hippocampus (Bender et al., 2021). In transgenic mice over-expressing α-synuclein, the accumulation of p-α-syn in the hippocampus resulted in an increase in early stage neural progenitor cells, though further studies have to be conducted to determine the fate of these cells (Bender et al., 2021). These results reveal a link between SIRT3 levels and age in humans, but also in PD pathology which is a disease where the primary risk factor is aging.

### In the PFF rat model, elevation of SIRT3 reduces α-synuclein aggregation

In clinical PD, α-synuclein aggregates spread across vulnerable brain regions (Abdelmotilib et al., 2017; Paumier et al., 2015; Rey et al., 2018). Since PFF α-synuclein models of PD are the only models that replicate the spread of α-synuclein pathology, we used it to further validate whether SIRT3 has disease-modifying effects.

In the control group infused with monomeric α-synuclein (mono+EV) and empty vector (EV) as a control for recombinant AAV1.SIRT3-myc, p-α-syn immunoreactivity was almost completely absent. This was expected since monomericα-synuclein does not aggregate. Following unilateral, intra-nigral infusion of PFFs, significantly higher levels of p-α-syn were observed in the SNc ipsilateral to the PFF infusion site compared to the mono+EV group. In the PFF+EV group, p-α-syn immunoreactivity was also present in the striatum, amygdala, hippocampus, motor cortex and olfactory tubercle. In human subjects with PD, these brain regions have also been shown to also exhibit α-synuclein pathology (Braak et al., 2004; Braak et al., 2003). Thus, although intra-nigral infusion of PFFs did not induce behavioural effects, it is a useful model for studying PD as it replicates the aggregation and spread of α-synuclein pathology in PD.

Phosphorylated α-synuclein was also present in the hemisphere contralateral to the PFF infusion, although to a much lower extent. This has also been shown previously, and is to be expected, since the two hemispheres are highly connected (Abdelmotilib et al., 2017; Rey et al., 2018; Volpicelli-Daley et al., 2014). Interestingly, in the hemisphere ipsilateral to PFF injection, levels of p-α-syn in the hippocampus and cortex were comparable to those in the SNc. Alpha-synuclein aggregates may spread via synaptic connections from the substantia nigra to the hippocampus, for example via the nigro-striatal, then ventro-striatal–hippocampal efferents (Sun et al., 2022; Zhang et al., 2017). It is also possible that immunoreactivity in the hippocampus and cortex may also be a result of fibrils diffusing from the needle tract during stereotactic surgery. Spread of p-α-syn from the needle tracts has also been shown previously (Harms et al., 2017; Lai et al., 2021).

AAV1.SIRT3-myc transduction significantly decreased p-α-syn abundance in the SNc. Additional replicates are required to deduce conclusively whether SIRT3 over-expression might also reduce p-α-syn in other brain regions. As the viral transduction of SIRT3 was localised to the SNc in conjunction with the PFF injection site, the reduction in p-α-syn aggregation in the SNc may reduce the seeding efficiency and thus the spread of pathological p-α-syn to interconnected brain regions.

In this study, infusion of PFFs had no significant effect on forelimb asymmetry at any timepoint, possibly due to the spread of PFFs to the contralateral hemisphere. While degeneration of dopaminergic neurons was not observed in the contralateral SNc, the prolific spread of PFFs to the contralateral hemisphere, in the striatum and motor cortex for example is likely to impact motor function. This pattern of spread of PFFs from the ipsilateral to contralateral side has been shown previously, particularly following injection of mouse PFFs, which appear to be more aggressive than human PFFs (Fares et al., 2016; Luk et al., 2012b; Masuda-Suzukake et al., 2013; Rey et al., 2016; Van Den Berge et al., 2021). While behavioural impairment was not observed in our PFF rat model, this is not unusual for the PFF model. While some groups have reported behavioural impairment in both mice and rats following administration of PFFs (Burtscher et al., 2020; Izco et al., 2021; Luk et al., 2016; Luk et al., 2012a; Masuda-Suzukake et al., 2014), others have observed no motor dysfunction (Luk et al., 2012a; Masuda-Suzukake et al., 2013; Rey et al., 2016). For the majority of these studies, PFFs were injected into the striatum for time periods of less than six months (Burtscher et al., 2020; Izco et al., 2021; Luk et al., 2016; Luk et al., 2012a; Masuda-Suzukake et al., 2014), which likely impacts model development, including the extent to which α-synuclein pathology spreads to the contralateral hemisphere. Additionally, in some studies behavioural impairment only occurs when PFFs are administered into transgenic animals (Ayers et al., 2018; Bieri et al., 2019; Gentzel et al., 2021) or with virally over-expressed α-synuclein (Espa et al., 2019), where mutant or wild-type α-synuclein levels in the brain are elevated well above amounts found in subjects with PD. Since no behavioural changes occurred in our PFF model, we were not able to assess the effect of AAV1.SIRT3-myc transduction on motor function in this study.

All groups that received nigral infusion of PFFs showed significant loss of dopaminergic neurons in the ipsilateral SNc. We believe the inability of SIRT3 to prevent degeneration of dopaminergic neurons is due to the aggressive cell loss induced by mouse PFFs compared to human PFFs when administered in rodents, as has previously been described (Luk et al., 2016). The reduction in p-α-syn indicates an effect of SIRT3 over-expression on α-synuclein aggregation. However, SIRT3 over-expression was not sufficient to completely protect the dopaminergic neurons in the SNc over a period of 6 months. While SIRT3 has been shown to interact with α-synuclein, the mechanism for SIRT3 mediated decrease in p-α-syn has yet to be elucidated. However, given that we and others have shown that elevation of SIRT3 stabilises mitochondria respiration and oxidative stress (Gleave et al., 2017), we predict the ability of SIRT3 to reduce α-synuclein aggregation is likely due to restoration of mitochondrial homeostasis and reducing oxidative stress. Given that oxidative stress results in enhanced synuclein aggregation, through interactions with dopamine quinones and other reactive oxygen species, reducing oxidative stress may reduce synuclein aggregation. Additionally, SIRT3-mediated stabilisation of mitochondria metabolisms may provide dopaminergic neurons with an increased capacity restore proteostasis allowing α-synuclein fibrils to be broken down.

Our findings that augmenting SIRT3 has disease-modifying effects might be consistent and related to findings of a recent Phase I clinical trials showing mild symptomatic improvements following oral administration of nicotinamide riboside (NR)(Brakedal et al., 2022)https://clinicaltrials.gov/ct2/show/NCT05589766). NR is the precursor to NAD+, and SIRT3 is both NAD+-dependent and activates pathways involved in replenishing NAD+ (Fulco et al., 2008; Xie et al., 2017). Importantly, many of the beneficial effects of NAD+ are mediated through SIRT3 activation, and SIRT3 declines with age. Thus, while increasing NAD+ levels through elevation of NR is an attractive target, it is possible that as the disease progresses, there will be insufficient SIRT3 to mediate the beneficial effects of NAD+. Furthermore, elevation of SIRT3 as a gene therapy may be attractive as a disease-modifying therapy because of the low frequency of administration (e.g., every 6 – 12 months).

## Conclusion

This study is the first to link age of the brain with reduced SIRT3 in control human subjects. Our findings also strongly suggest that SIRT3 is directly involved with the pathological development of clinical PD. In clinical PD, many of the known mitochondrial dysfunctions are normally regulated by SIRT3, such as oxidative stress, reduced complex I activity, and dysregulation of mitochondrial proteostasis. Furthermore, SIRT3 has been shown to declines in aging skeletal muscle, and age is the number one risk factor of PD. Thus, reduced SIRT3 may be a key driver of PD pathology. The ability of ectopically expressed SIRT3 to reduce the aggregation of α-synuclein in the SNc, combined with our previous studies showing neurorestorative effects *in vivo* (Gleave et al., 2017) suggest that AAV1.SIRT3-myc could serve as a disease-modifying gene therapy. By simultaneously targeting α-synuclein aggregation and mitochondrial dysfunction in subjects with PD, SIRT3 over-expression provides a two-pronged approach to combatting the neurodegenerative process.

## Methods

### Animal care

Male Sprague Dawley rats (275-325g) (Charles River, Canada) were housed in pairs in a temperature-controlled environment with a 12-hour light/dark cycle. Animals had free access to food and water, except 12 hours before behavioural assessment, as this has been shown to improve performance in the cylinder test All animal studies were performed in accordance with the Canadian Council on Animal Care and the Local Animal Care Committee at the University of Toronto Ethical Guidelines.

### Intra-nigral infusion of AAV1.SIRT3-myc

SIRT3-myc transduction was achieved using a recombinant adeno-associated virus serotype 1 (AAV1). AAV1 has previously been shown to be the most efficiently transduced natural serotype in neurons (Burger et al., 2004; Taymans et al., 2007). Also, we have previously optimised AAV1.SIRT3-myc. and shown it has disease-modifying effects in the mutant A53T α-synuclein rat model of PD (Gleave et al., 2017). AAV1.SIRT3-myc was infused into the SNc using stereotaxic surgery (Frame: Stoelting Co., USA, 1 cc glass syringe: Hamilton Company, USA), and microinjector (Stoelting Co., USA). AAV1.SIRT3-myc (University of Pennsylvania Vector Core, USA) was infused into the right SNc (coordinates: AP −5.2mm, ML −2.0 mm, DV −7.5 mm from Bregma according the atlas of Paxinos at an infusion rate of 0.20 μL/min (total volume 2 μL) using a stock solution with a concentration of 2.59 x 10^11^ GC/mL.

### Generation of a mis-folded α-synuclein rat model of PD using pre-formed fibrils

Pre-formed mis-folded fibrils of mouse α-synuclein (PFFs) were generated in the laboratory of Dr. Laura A. Volpicelli-Daley (University of Alabama at Birmingham, USA) following the protocol as described previously (Luk et al., 2012a; Luk et al., 2012b; Luk et al., 2009). PFFs were stored at −80 °C immediately after generation and thawed on ice prior to use. Activation of 100 μL aliquots of PFFs was achieved using a probe tip sonicator with 1 second on/off bursts for 1 minute. Successful activation as confirmed by the shortening of the PFFs from approximately 200nm to 50nm was visualised using transmission electron microscopy. Monomeric α-synuclein served as the control for PFFs, was centrifuged at 21700*g* (4 °C) for 1 hour to remove small aggregates from the supernatant. Both activated PFFs and monomeric α-synuclein were stored at room temperature (RT) during stereotaxic surgeries. PFFs or monomeric α-synuclein (5 ng/μL) were infused into the right SN (coordinates: AP −5.3 mm, ML −2.2 mm, DV −7.3 mm from Bregma according to the rat brain atlas using stereotaxic surgery at an infusion rate of 0.25 μL/min (4 μL total volume) using a 1 cc glass syringe (Hamilton Company, USA) and microinjector (Stoelting Co., USA).

### Cylinder test to assess forelimb asymmetry

Given that AAV1.SIRT3-myc and PFFs were infused into the right hemisphere, we hypothesised that any behavioural effects caused by transduction of AAV1.SIRT3-myc or PFFs would be unilateral, and so cause forelimb asymmetry. Thus, behavioural assessment of the effects of PFF infusion and SIRT3 transduction were measured using the cylinder test, which measures forelimb asymmetry. Rats were assessed at the beginning of the light cycle after being food deprived for 12 hours during the dark cycle. Testing involved rats being placed individually into a clear glass cylinder (dimensions: 19.2 cm D x 30.7 cm H) surrounded by 2 mirrors placed in such a way that all angles of the cylinder could be recorded on camera. Each rat underwent 3 trials with the video capturing 25 total forelimb contacts against the walls of the cylinder, recording for no longer than 6 minutes. Forelimb asymmetry was then analysed using a single blinded technique, where the first 20 forelimb placements were assessed as ipsilateral or contralateral paw placements, or both relative to the right hemisphere. The following equation was used to calculate forelimb asymmetry:

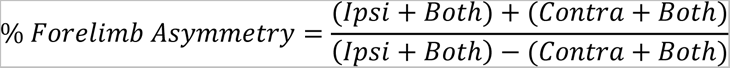

### Tissue collection and preparation

Six months following PFF-infusion, rats were euthanised using isoflurane (3-5%) and cardiac perfusion performed with ice cold PBS. Brains were removed and either fixed in ice cold 4% PFA overnight at 4 °C or dissected to obtain the striatum which was immediately flash frozen. Fixed brains were then washed 3 times in ice cold PBS before being incubated in 30% sucrose solution until the brains sank. Cryosectioning was performed to obtain 40μm sections of brain tissue and stored in cryoprotectant at −20 °C until further processing occurred.

### Immunohistochemistry/immunofluorescence

For immunohistochemistry, brain sections were removed from cryoprotectant and washed 3 times for 10 minutes in PBS-0.1% Triton X (PBS-T), followed by a 15minute incubation in 3% hydrogen peroxide. Sections were then washed 3 times for 5 minutes in PBS-T before blocking for one hour in 5% normal goat serum (NGS) in PBS-T. Following the block, sections were incubated overnight at 4°C in primary antibodies. Sections were then washed with PBS-T 3 times for 5 minutes, and incubated 2 hours at RT in secondary antibodies diluted in 1% NGS in PBS-T. After 3 5 minute washes with PBS-T, the sections were incubated in ABC Elite reagent (Vector Laboratories, USA) for 1 hour at RT following the manufacturer’s protocol. Sections were then rinsed 3 times for 5 minutes with PBS, then stained with DAB (Vector Laboratories, USA) following the manufacturer’s protocol. Sections requiring labelling for TH following NeuN received 3 5minute washes in PBS-T, followed by an overnight incubation at RT with primary antibody in 1% NGS in PBS-T. The next day, sections were washed 3 times for 5 minutes in Tris buffered saline-0.1% Tween 20 (TBS-T) and incubated in secondary antibodies in 1% NGS in TBS-T for two hours at RT, which was then washed off 3 times for 5 minutes in TBS-T. Staining with Vector Blue substrate (Vector Laboratories, USA) was performed according to the manufacturer’s protocol. Stained sections were then washed 3 times for 5 minutes in PBS, mounted on glass microscope slides and allowed to air dry overnight protected from light. Slides were then submerged in ddH_2_O for three minutes, 70% EtOH for one minute, 95% EtOH for one minute, 100% EtOH for one minute, Histoclear twice for 3 minutes, then mounted using Vectamount (Vector Laboratories, USA).

For immunofluorescence, brain sections were removed from cryoprotectant and washed 3 times for 10 minutes in PBS-T, followed by blocking in 5% NGS in PBS-T for 1 hour. Sections were then incubated overnight at 4 °C in 1% NGS in PBS-T containing primary antibodies. The next day sections were washed with PBS-T 3 times for 5 minutes and incubated with secondary antibody in 2% NGS in PBS-T for 1 hour at RT. Following 3 5minute washes in PBS, sections were mounted onto glass microscope slides. After drying, sections were rehydrated and washed with ddH_2_O for 2 minutes and mounted using Mounting Medium with DAPI (Abcam, USA), and edges sealed with clear enamel once the mounting medium set.

### Quantitative stereological analysis

To estimate the number of dopaminergic neuron population in the SNc, quantitative stereology was performed. Serial sections of SNc (1:6) 40μm in thickness labelled for NeuN and TH using immunohistochemistry were analysed using Stereo Investigator software on a Zeiss Imager M2 microscope (Zeiss, Germany) to estimate the number weighted section thickness dopaminergic neuron population (NeuN/TH positive cells). Following the quantification of NeuN/TH positive cells in the SNc (with the region being user-defined with a 5x objective lens), the population of dopaminergic neurons was calculated by the software. The outlined region was divided by the software into 120μm x 90μm counting grids, with a 60μm x 60μm counting frame surrounded by 2μm of guard space. Section thickness was evaluated at each sampling site containing at least 1 NeuN/TH positive cell, which were analysed sequentially with an 100x oil immersion lens.

### Quantitative phosphorylated Ser129 α-synuclein analysis

Brain sections containing the SNc, striatum, hippocampus, amygdala, motor cortex, and olfactory tubercule were imaged using a Leica SP8 confocal microscope (Leica Microsystems, Germany) following immunofluorescence labeling for p-α-syn. Tile scans containing the regions were imaged with a 20x objective lens, and then traced using ImageJ. The amount of p-α-syn in each region was determined by calculating the percentage area (pixels) with p-α-syn positive signal divided by the total area of the region.

### High-performance liquid chromatography

To quantify dopamine and its metabolites, HPLC was performed on flash frozen samples of rat brain striatum by the Neurochemistry Core lab (Vanderbilt University, USA). Homogenates were centrifuged and the resulting supernatant analysed using an Antec Decade II electrochemical detector operated at 33*^°^*C and C_18_HPLC column (100 x 4.60 mm). Biogenic amines were eluted with a mobile phase (89.5% 0.1 M TCA, 0.01 M sodium acetate, 0.0001 EDTA, and 10.5% methanol, pH 3.8).

### Human tissue SDS-PAGE and Western Blot analysis

Human brain tissue (hippocampus and SNc) from Parkinson’s disease and control subjects was obtained from the Douglas Brain Bank. Sections of hippocampus and SN were weighed, and lysis buffer containing 50 mM Tris pH 8.0, 1% NP40, 150 mM NaCl, 1 mM EDTA, 1 mM PMSF, 1 μg/mL Aprotinin, 1 μg/mL Leupeptin, 2 mM Na_3_VO_4_ and supplemented with protease inhibitor cocktail tablets was added at a 1:1 weight to volume ratio. Tissue was homogenized using a 1mL glass Dounce homogenizer for approximately 35 strokes, then centrifuged at 10000g for 10 minutes (4*°*C). The resultant supernatant containing the proteins of interest was collected, and protein concentration determined using a Lowry protein assay. Protein samples (15ng) diluted with TBS (2.42 g Tris, 8 g NaCl, pH7.6) and Laemmli (4X stock diluted to 1X during sample preparation) loaded into a 0.8% acrylamide SDS-PAGE gel and ran through the stacking layer at 65 V and the separating layer at 125 V. Proteins were transferred onto a nitrocellulose membrane (0.2 A, 1 hour), and blocked one hour in 5% skim milk powder (SMP) in TBS-0.05% Tween 20, then rinsed 3 times using TSB-Tween 20 before incubating overnight with primary antibodies in 2% albumin TSB-Tween 20 (4°C). Then the membrane was washed 3 times for 10 minutes in TBS-Tween 20 then incubated for two hours in 1% SMP TBS-Tween 20 with secondary antibodies. Membranes were developed after 3 10 minute washes in TBS-Tween 20 using Amersham ECL Developing Reagent and imaged using the Bio-Rad ChemiDoc XRS+. Membranes were then stripped for 15 minutes in Restore Western Blot Stripping Buffer and washed 3 times for 10 minutes in TSB-Tween 20 before relabeling from the blocking step as described above.

### Statistics

Statistical analysis was performed using Prism 9.3 software (GraphPad Software Inc.). Data are presented as mean ± SEM. For behavioural assessment, one-way ANOVA followed by Tukey’s multiple-comparison test was performed while quantitative analysis of dopaminergic neurons, and striatal dopamine and its metabolites were tested using a one-way ANOVA followed by Bonferroni’s multiple-comparison test. Abundance of p-α-syn in the brain was tested using a two-way ANOVA followed by Bonferroni’s multiple-comparison test. Comparisons between control and PD subjects Western Blot protein expressions were performed using Mann-Whitney test (non-parametric). Regression analysis of Western Blot data was tested using the Pearson Correlation test. Significance was indicated by p<0.05. Sample sizes are provided in the figure legends.

### Study Approval

All animal procedures and experiments were conducted under the approval of the Local Animal Care Committee at the University of Toronto in accordance with the regulations and guidelines set by the Canadian Council on Animal Care. Procedures and experiments using human tissue were conducted under the approval of the Human Research Ethics Unit at the University of Toronto. Written consent was provided by subjects to the Douglas Brain Bank where the human tissue samples were obtained.

## Author Contributions

Experiments were designed by DT^1^, ARI, and JEN. Experiments were performed by DT^1^ and ARI. Data was acquired by DT^1^, ARI, HB, JEAV, and RL, and analysis was performed by DT^1^, ARI, HB, JEAV, RL, HP, KR, AS, VG, and NG. PFFs were provided by DT^3^ and LV. The manuscript was written by DT^1^ and JEN.

